# Comparative population genomics of manta rays has global implications for management

**DOI:** 10.1101/2023.06.19.545572

**Authors:** Emily Humble, Jane Hosegood, Gary Carvalho, Mark de Bruyn, Simon Creer, Guy M W Stevens, Amelia Armstrong, Ramon Bonfil, Mark Deakos, Daniel Fernando, Niv Froman, Lauren R Peel, Stephen Pollett, Alessandro Ponzo, Allan L Poquita, Joshua D Stewart, Sabine Wintner, Rob Ogden

## Abstract

Understanding population connectivity and genetic diversity is of fundamental importance to conservation. However, in globally threatened marine megafauna, challenges remain due to their elusive nature and wide-ranging distributions. As overexploitation continues to threaten biodiversity across the globe, such knowledge gaps compromise both the suitability and effectiveness of management actions. Here, we use a comparative framework to investigate genetic differentiation and diversity of manta rays, one of the most iconic yet vulnerable groups of elasmobranchs on the planet. Despite their recent divergence, we show how oceanic manta rays (*Mobula birostris*) display significantly higher heterozygosity than reef manta rays (*Mobula alfredi*) and that *M. birostris* populations display higher connectivity worldwide. Through inferring modes of colonisation, we reveal how both contemporary and historical forces have likely influenced these patterns, with important implications for population management. Our findings highlight the potential for fisheries to disrupt population dynamics at both local and global scales and therefore have direct relevance for international conservation of marine species.

## Introduction

Understanding the extent to which populations are connected is key to exploring population dynamics, predicting extinction risk and informing conservation management (Hanski & Gilpin, 1991; Lowe & Allendorf, 2010; Mills & Allendorf, 1996). In species with isolated populations characterised by limited dispersal, the risk of extirpation from local depletion is high (Reed, 2004). In such cases, local and regional scale management will be most appropriate for preventing and reversing population declines (Palumbi, 2003). In contrast, species with high rates of gene flow are potentially demographically and genetically more resilient to extrinsic factors (Lowe & Allendorf, 2010; Pascual et al., 2017). However, in order to maintain connectivity and mitigate genetic diversity loss in these taxa, management measures must be coordinated and encompass migratory corridors. As overexploitation and habitat destruction threaten to disrupt population dynamics at a global scale, characterising genetic variation and connectivity has become more important than ever before (Funk et al., 2012; Kardos et al., 2021; Palsbøll et al., 2007).

In widely distributed marine species with high dispersal potential, genetic differentiation is often found to be subtle or non-existent (Palumbi, 2003; Waples, 1998; Ward et al., 1994). Such patterns can arise from a range of mechanisms – from high contemporary gene flow to recent divergence of historically large populations (Palumbi, 2003; Waples et al., 2008; Waples & Gaggiotti, 2006) – and can therefore be difficult to interpret. The latter scenario reflects a disconnect between demographic and genetic connectivity and has important implications for species resilience (Bailleul et al., 2018; Lowe & Allendorf, 2010; Waples, 1998). This is because populations that appear genetically connected may not operate as single demographic units, making them more vulnerable to overexploitation. High-resolution SNP datasets go some way to addressing this problem by providing greater power to detect subtle differences at both neutral and adaptive loci (Gagnaire et al., 2015; Hauser & Carvalho, 2008). However, since population genetic differentiation can be affected by past, as well as contemporary patterns, parallel inference of historical relationships and genetic variation can allow the relative contribution of historical processes to be explicitly evaluated (Foote & Morin, 2016; Liu et al., 2022; Louis et al., 2021). Furthermore, when carried out within a comparative framework, such an approach can provide powerful insights into the drivers of population divergence and therefore improve recommendations for conservation management (Gagnaire, 2020).

Manta rays are large, mobile elasmobranchs inhabiting tropical and sub-tropical oceans (Couturier et al., 2012) (Figure 1A, C) and provide an excellent opportunity to evaluate the genomic consequences of historical and contemporary population processes within a comparative framework. They comprise two described species estimated to have diverged less than 0.5 Mya as a result of distinct habitat preferences (Kashiwagi et al., 2012). The reef manta ray (*Mobula alfredi*) frequents near-shore tropical reef environments, such as coral atolls and barrier reefs (Kashiwagi *et al*. 2011), with a high degree of residency (Deakos *et al*. 2011; Jaine *et al*. 2014; Braun *et al*. 2015; Setyawan *et al*. 2018; Peel *et al*. 2019; Knochel *et al*. 2022b; Germanov *et al*. 2022). In contrast, while the oceanic manta ray (*Mobula birostris*) also inhabits near-shore environments, it is often found ranging into sub-tropical habitats along continental coastlines and at oceanic islands, usually adjacent to productive deep-water upwellings (Kashiwagi *et al*. 2011; Andrzejaczek *et al*. 2021). As a result of these differences in habitat use, *M. alfredi* and *M. birostris* have long been considered to display marked differences in their migratory abilities and levels of gene flow. Yet, only a handful of long-distance movements have ever been recorded in *M. birostris* (Andrzejaczek et al., 2021; Knochel, Cochran, et al., 2022) alongside observations of site-fidelity (Cabral et al., 2023; Garzon et al., 2023; Gordon & Vierus, 2022), raising questions about the extent to which population structure and genetic variation may differ across species. To date, assessments of genetic differentiation in *M. alfredi* have focussed on local and regional patterns (Lassauce et al., 2022; Venables et al., 2021; Whitney et al., 2023) and we have little understanding of how genetic variation is distributed across the species’ range. In, *M. birostris*, the situation is even less clear, with studies reporting both widespread connectivity and population differentiation (Hosegood et al., 2020; López et al., 2022; Stewart et al., 2016). Critically, these differences and uncertainties exist against a background of ongoing global exploitation and uncertain implications for management.

**Figure 1.**
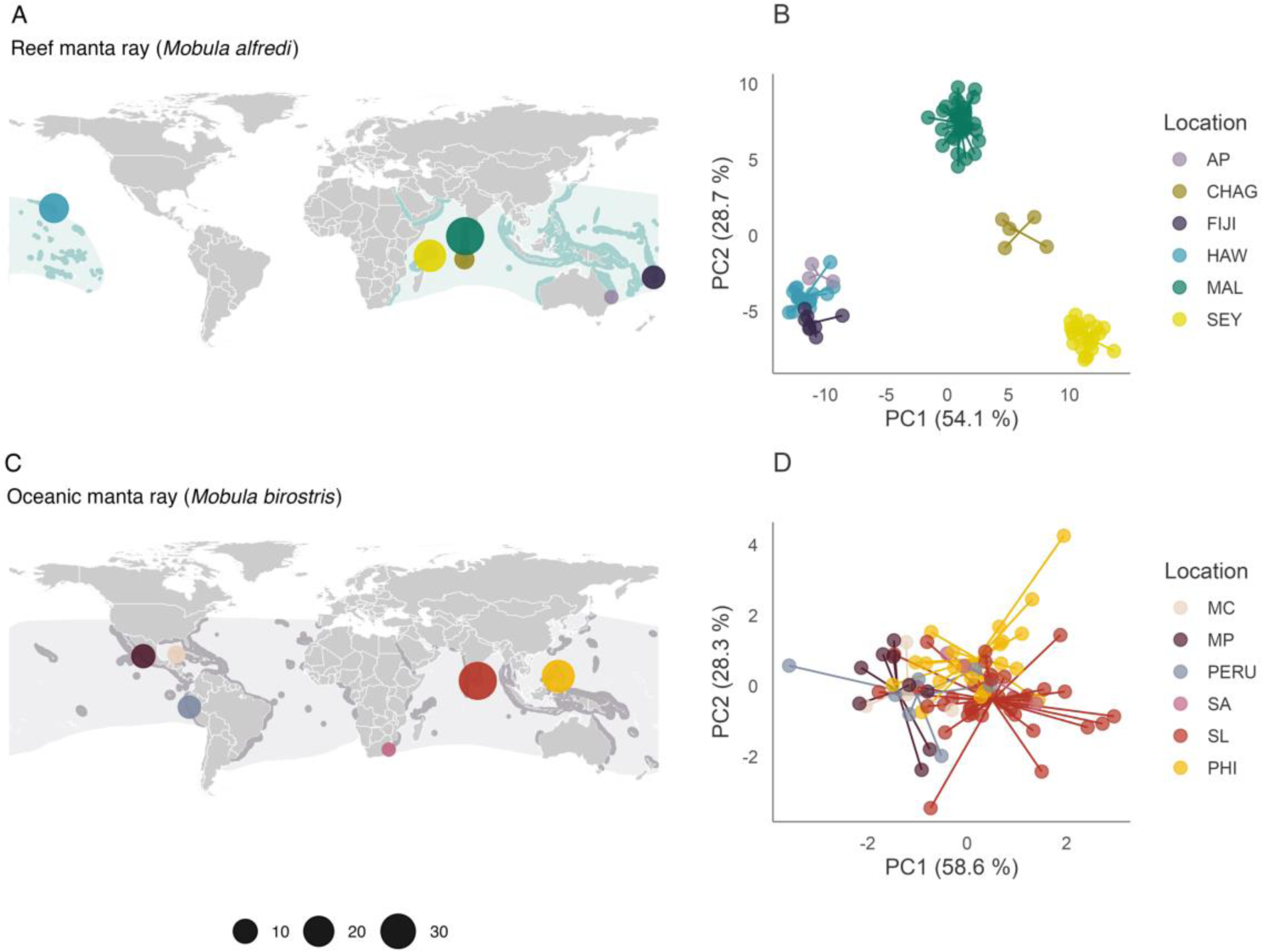
Contrasting patterns of population structure in manta rays. (A, C) Geographic distributions of (A) *Mobula alfredi* and (C) *Mobula birostris* visualised together with the locations of samples used in this study. Dark shaded distributions denote the confirmed species range and light shaded distributions denote the expected species range. Sampling location points are distinguished by colour and scaled by the number of samples. Further details are provided in the Supplementary Material. (B, D) Scatterplots showing individual variation in principal components (PC) one and two derived from discriminant analysis of principal components analysis for (B) *M. alfredi* and (D) *M. birostris* individuals. The amount of variance explained by each PC is shown in parentheses. Population abbreviations: AP = Australia Pacific, CHAG = Chagos, FIJI = Fiji, HAW = Hawaii, MAL = Maldives, SEY = Seychelles, MC = Mexico Caribbean, MP = Mexico Pacific, PERU = Peru, SA = South Africa, SL = Sri Lanka and PHI = the Philippines.

Targeted and incidental fisheries, driven in part by increasing demand for mobulid gill plates (Couturier et al., 2012; O’Malley et al., 2017), have led to widespread population declines in manta rays (Carpenter et al., 2023; Croll et al., 2016; Moazzam, 2018; Rohner et al., 2017; Ward-Paige et al., 2013). Currently, both species are managed through a patchwork of local, regional and international measures with varying levels of implementation and enforcement (Fernando & Stewart, 2021; Lawson et al., 2017; Lawson & Fordham, 2018). To determine the appropriateness of management measures and assess population vulnerability, a global assessment of management units is urgently required (Lawson et al., 2017; Stewart, Jaine, et al., 2018). Here, we undertake a comparative genomic analysis of manta ray populations from across their global distribution to investigate connectivity, genetic variation, and historical relationships with an aim to guide effective fisheries management and emphasise the value of genomic research for advancing knowledge of understudied elasmobranchs.

## Materials and Methods

### Sample collection

Tissue samples were opportunistically collected from 12 geographic locations to represent the global distribution of each species (Figure 1A, C). For *M. alfredi* (total *n* = 120), these originated from the Chagos Archipelago (n = 5), the Maldives (n = 48), Seychelles (n = 24), Australia Pacific (n = 4), Fiji (n = 9) and Hawaii (n = 30). For *M. birostris* (total *n* = 112), these originated from Sri Lanka (n = 43), the Philippines (n = 37), South Africa (n = 3), Mexico Caribbean (n = 4), Mexico Pacific (n = 13) and Peru (n = 12). Samples from Mexico Caribbean, where a third putative manta ray species occurs in sympatry (*Mobula cf. birostris,* Hinojosa-Alvarez *et al*. 2016; Hosegood *et al*. 2020), were visually and genetically confirmed as *M. birostris*. For both species, samples were collected from a combination of live animals and fisheries specimens (See Supplementary Files for further information).

### DNA extraction and ddRAD sequencing

Genomic DNA was extracted using the Qiagen DNeasy Blood and Tissue Kit and quantified using a Qubit 3.0 Broad Range Assay. Double digest restriction-site associated DNA (ddRAD) libraries were prepared following the Peterson *et al*. (2012) protocol with modifications described in Palaiokostas *et al*. (2015) and were 125 bp paired-end sequenced on an Illumina HiSeq. Sequencing reads from both species were assessed for quality using FastQC and processed together using the Stacks v2.54 *de novo* assembly pipeline (Catchen et al., 2013). The three main assembly parameters were chosen following the optimisation procedure outlined in Rochette and Catchen (2017) (Figure S1 and S2). Initial quality filters were applied to the resulting genotypes before generating three high-quality datasets for use in downstream analysis: two species-specific datasets; and one dataset comprising both species. For the species-specific datasets, we extracted either *M. birostris* or *M. alfredi* individuals, removed individuals with high relatedness coefficients (Korneliussen & Moltke, 2015; Waples et al., 2019, Figure S3) and filtered out SNPs with a minor allele count of less than 3, a genotyping rate less than 90% and that were in linkage disequilibrium using PLINK. For the dataset comprising both species, we first removed closely related individuals and then filtered out SNPs with a minor allele count of less than 3 and a genotyping rate less than 90%. Population genetic summary statistics were calculated using the R package diversity (Keenan et al., 2013, Table S1). See Supplementary Material for further information on library preparation, read processing and SNP and individual filtering.

### Population structure

To investigate population structure we used the species-specific datasets and three complementary approaches. First, we carried out a discriminant analysis of principal components (DAPC) using the R package adegenet (Jombart, 2008). This approach initially transforms the SNP data using a principal components analysis (PCA) and then performs a discriminant analysis on the retained PCs. This serves to maximise discrimination of individuals between groups while minimising variation within (Jombart et al., 2010). Following the recommendations outlined in Thia (2023), the number of PCs retained as predictors was determined based on the *K–1* criterion, where *K* is equal to the number of effective populations. For *M. alfredi*, this was set to 5, under the assumption that each sample site reflects a separate population. For *M. birostris*, this was set to 4 under the assumption that Mexico Pacific and Peru may represent a single population given their close geographic proximity. Second, we estimated admixture proportions for the individuals in each dataset using ADMIXTURE. Admixture runs were performed for ancestry clusters ranging from *K* = 1– 8, with 10 runs for each *K*. The optimal *K* was identified based on the lowest cross-validation error. The runs with the highest likelihood were visualised. Third, we estimated pairwise genetic differentiation between populations within each species using the Weir and Cockerham *F*_ST_ value (Weir & Cockerham, 1984) calculated in the R package dartR (Gruber et al., 2018). Confidence intervals and *p-*values were estimated based on bootstrap resampling of individuals within each population 1000 times. *Mobula alfredi* samples from Australia Pacific and *M. birostris* samples from South Africa were excluded from this analysis due to low sample sizes.

### Isolation by distance

To investigate patterns of isolation by distance, we examined the relationship between genetic and geographic distance between all pairs of populations in each species. Genetic distances were based on pairwise *F*_ST_ estimates calculated above. Geographic distances were determined based on a least-cost path analysis implemented using the R package marmap (Pante & Simon-Bouhet, 2013) with a minimum depth constraint of −10 metres in order to prevent paths overland. The significance of associations between genetic and geographic distance matrices was inferred using distance-based Moran’s eigenvector maps (dbMEM) by redundancy analysis (RDA, Legendre et al., 2015). For this, geographical distances were transformed into dbMEMs using the R package adespatial, and genetic distances were decomposed into principal components using the R function prcomp. RDA was then performed using the R package vegan, with significance tested using 1000 permutations.

### Contemporary gene flow

To infer the strength and directionality of contemporary gene flow between populations we used the program BA3-SNPs BayesAss v1.1 (Mussmann et al., 2019) which estimates the proportion of immigrants in a given population using Bayesian inference. This analysis was restricted to *M. alfredi* as it assumes low levels of connectivity and imposes an upper-bound on the proportion of non-migrants in a population. We first performed initial runs of BayesAss to determine optimal mixing parameters (dM = migration rate, dA = allele frequency and dF = inbreeding coefficient) using the autotune function in BA3-SNPs. We than ran BayesAss-3 with 10,000,000 iterations, a burn-in of 1,000,000 and a sampling interval of 1000. Mixing parameters were set to dM = 0.21, dA = 0.44 and dF = 0.08. Results were averaged across five replicate runs and migration rates were considered significant if 95% credible sets (mean migration rate ± 1.96 x mean standard deviation) did not overlap zero. Chain convergence was assessed, and migration rates visualised using R (Figure S4).

### Historical relationships among populations

To explore historical relationships among populations of *M. alfredi* and *M. birostris* we used the program TreeMix (Pritchard et al., 2000). TreeMix uses population allele frequencies to estimate a bifurcating maximum likelihood tree with which to infer historical population splits, admixture events and the degree of genetic drift. We first supplemented the *M. alfredi* dataset with one randomly selected *M. birostris* individual and the *M. birostris* dataset with one randomly selected *M. alfredi* to act as outgroups when rooting the trees. Both datasets were then filtered for linkage, a minor allele count of less than 3, genotyping rate of less than 90% and related individuals using PLINK v1.9 (Purcell et al., 2007). Allele frequencies for each population were then calculated using the –freq and –within arguments in PLINK. For both the *M. birostris* and *M. alfredi* datasets we then performed 10 initial runs of TreeMix for each migration event (M) ranging from 0 to 10. The number of migration edges that explained 99.8% of the variance was selected as the best model for each species (*M. birostris:* M= 0; *M. alfredi:* M = 2, Figure S5). We then re-ran TreeMix 100 times using the optimal number of migration edges. Consensus trees and bootstrap values were estimated and visualised using code modified from the BITE R package (Milanesi et al., 2017).

### Genome-wide heterozygosity

To assess levels of genetic variation both within and between species we used the high-quality SNP dataset comprising both species. We then calculated multi-locus heterozygosity for each individual using the R package inbreedR (Stoffel et al., 2016).

## Results

High-quality SNPs were genotyped in 173 individuals from 12 locations representing the global distribution of each species (Figure 1A, C). The species-specific datasets contained a total of 1,553 SNPs in 91 *M. alfredi* individuals, and 6,278 SNPs in 82 *M. birostris* individuals, while the full dataset contained a total of 15,312 SNPs called across both species. See Materials and Methods and Supplementary Information for details.

### Contrasting patterns of population structure at a global scale

To investigate population differentiation within each species we used four complementary approaches: discriminant analysis of principal components (DAPC), admixture, pairwise *F*_ST_ and isolation by distance analysis. In *M. alfredi*, all methods supported the presence of strong population structure at both global and regional scales. Populations inhabiting different ocean basins displayed the highest degree of differentiation in the DAPC, with Pacific and Indian Ocean populations forming distinct clusters along PC1 (Figure 1B). Regional differentiation was also detected, with Seychelles, Chagos and the Maldives clustering apart along PC2, and Hawaii separating from Australia Pacific and Fiji along PC3 (Figure 1B and Figure S6A). These patterns were reinforced in the admixture analysis which highlighted two major ancestral source populations, inferred an optimal value of *K* = 4 and resolved hierarchical structure up to *K* = 7 (Figure S7 and Figure S8A). Interestingly, only weak separation was observed between Australia Pacific and Fiji, however, this pattern may be confounded by the small sample size of the former, which can lead to spurious merging of distinct populations (Puechmaille, 2016). Pairwise *F*_ST_ estimates between ocean basins were on average over two times higher than those within (mean pairwise *F*_ST_ between ocean basins = 0.30, mean pairwise *F*_ST_ within ocean basins = 0.13, Figure 2A) yet all population comparisons were found to be significant (Figure S9A, mean = 0.23, min = 0.08, max = 0.43). Finally, we detected a significant relationship between pairwise *F*_ST_ and geographic distance (adjusted *R^2^*= 0.65, *P* = 0.03) indicating an effect of isolation by distance (Figure 2B).

**Figure 2.**
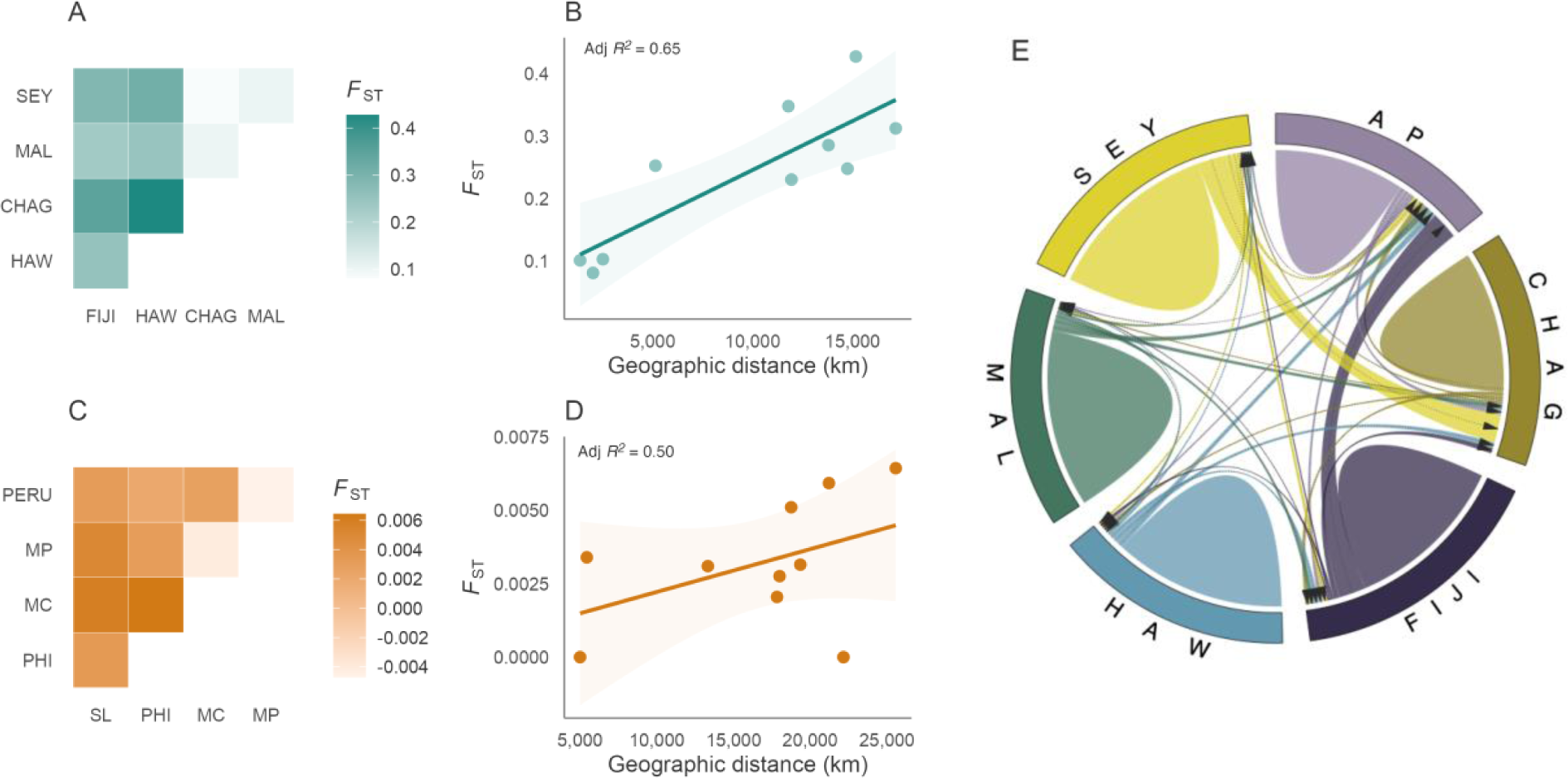
Population genetic differentiation, isolation by distance and contemporary migration in manta rays. (A, C) Pairwise *F*ST estimates between sampling locations for (A) *M. alfredi* and (C) *M. birostris*. Samples from Australia Pacific and South Africa were excluded from this analysis due to low sample sizes. (B, D) Relationship between genetic (*F*ST) and geographic distance as calculated by least-cost path analysis for all pairwise population comparisons in (B) *M. alfredi* and (D) *M. birostris*. Solid lines and shaded areas reflect the regression slopes and standard errors respectively based on a linear model. Samples from Australia Pacific and South Africa were excluded from this analysis due to low sample sizes. (E) Contemporary gene flow estimates between populations of *M. alfredi*. The direction of each arrow represents the direction of gene flow, and the width of each ribbon reflects the relative amount of gene flow. Population abbreviations: AP = Australia Pacific, CHAG = Chagos, FIJI = Fiji, HAW = Hawaii, MAL = Maldives, SEY = Seychelles, MC = Mexico Caribbean, MP = Mexico Pacific, PERU = Peru, SL = Sri Lanka and PHI = the Philippines.

In stark contrast, *M. birostris* displayed little evidence for strong population structure across all methods. Individuals from different ocean basins clustered closely together along each axis in the DAPC (Figure 1D and Figure S6B). Admixture identified *K* = 1 as the optimal number of clusters, with increasing values of *K* merely introducing additional mixing (Figure S7 and Figure S8B). Pairwise *F*_ST_ estimates were two-fold lower than in *M. alfredi*, with no pairwise comparison falling above 0.007 (mean = 0.002, min = −0.005, max = 0.006, Figure 2C). Nevertheless, despite these broad patterns, several lines of evidence indicate the presence of subtle geographic differentiation in this species. First, individuals from Mexico Pacific, Peru and Mexico Caribbean clustered separately from those sampled in South Africa, Sri Lanka, and the Philippines along PC1 (Figure 1D). Second, despite pairwise *F*_ST_ estimates being low, comparisons between Eastern-Pacific and Indo-Pacific populations, and between Sri Lanka and the Philippines were statistically significant (Figure S9B). Small *F*_ST_ values are expected when minor allele frequencies are low and therefore do not necessarily reflect an absence of differentiation (Jakobsson et al., 2013). Furthermore, it is possible that overall levels of population structure were underestimated due to the small sample size of two of our six populations (Puechmaille, 2016). Finally, while no significant relationship was observed between pairwise *F*_ST_ and geographic distance (adjusted *R^2^*= 0.50, *P* = 0.10), there was a tendency for populations separated by greater distances to display higher differentiation (Figure 2D).

### Contemporary gene flow

To characterise the strength and direction of gene flow between populations we used the program BA3-SNPs (Mussmann et al., 2019) to estimate recent migration. As this method assumes low levels of connectivity and imposes an upper-bound on the proportion of non-migrants in a population, this analysis was restricted to *M. alfredi*. As expected, contemporary gene flow was low (Figure 2E); the average migration rate between populations, measured as the estimated number of migrants per generation, was 0.029 (min = 0.008, max = 0.15), with this figure falling to 0.018 (min = 0.008, max = 0.03) when considering gene flow between populations in different ocean basins. Migration into both Hawaii and the Maldives was lowest, indicating these populations are the most isolated of those sampled (Table S2). Migration rates were only deemed significant between Seychelles and Chagos (0.15) and between Fiji and Australia Pacific (0.15), in line with these populations being last to separate in the admixture analysis. These patterns highlight that while *M. alfredi* may have the propensity to travel over large distances, restricted movement likely dominates.

### Historical relationships among populations

To place patterns of genetic differentiation into a historical context, we investigated population origins and colonisation patterns using TreeMix (Pritchard et al., 2000). This program uses allele frequency data to infer patterns of population splits and admixture events through the construction of a maximum likelihood tree. In *M. alfredi*, internal branch lengths were relatively long, with an initial split clearly separating populations in the Indian and Pacific Oceans (Figure 3A). The Maldives and Australia Pacific were the first to separate within each locality and displayed the lowest levels of genetic drift overall. Hawaii was among the last populations to split and displayed the highest amount of drift, in line with its geographic isolation. The best supported model inferred two migration events (Figure S5A); one from the *M. alfredi* population in Seychelles into *M. birostris*, and one from Chagos into Hawaii. However, because not all geographic regions are represented in our data set, the true sources and sinks of these admixture events may originate from related ghost populations. In contrast to *M. alfredi*, the addition of migration events led to no substantial improvement in the model for *M. birostris* (Figure S5B) and therefore the tree without migration is presented here. Interestingly, internal branch lengths were considerably shorter in *M. birostris*, indicating rapid radiation from a shared ancestral source population (Figure 3B). External branch lengths were also short, consistent with larger populations displaying marginal drift and low divergence. Nevertheless, despite these patterns, some geographic signal was detected in the *M. birostris* tree, with individuals originating from the Eastern Pacific and the Caribbean (Peru, Mexico Caribbean, and Mexico Pacific) grouping separately from those originating from the Atlantic and Indo-Pacific (South Africa, the Philippines, and Sri Lanka), although bootstrap support was overall low.

**Figure 3.**
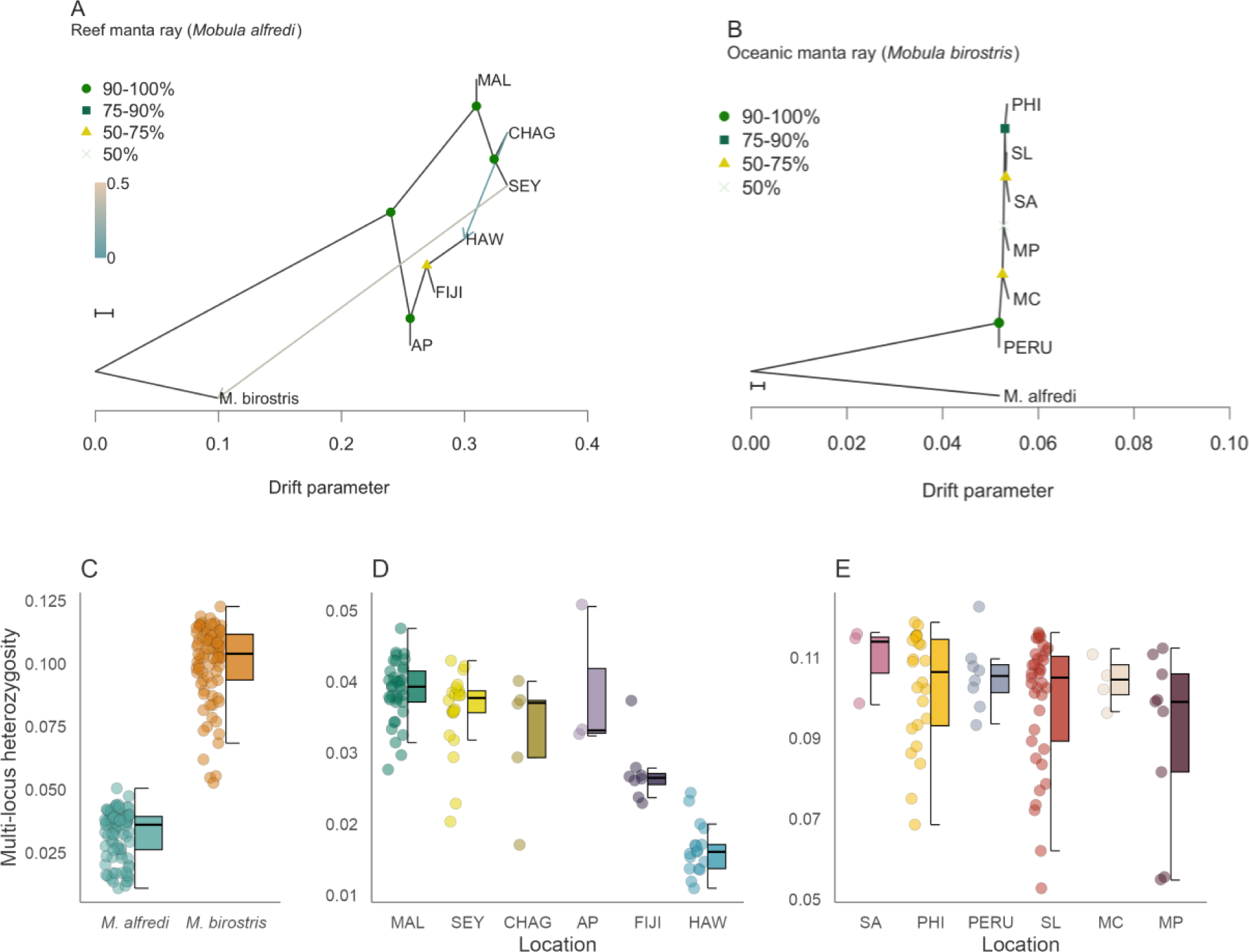
Historical relationships and heterozygosity in manta rays. (A–B) TreeMix maximum likelihood consensus tree displaying the historical relationships among (A) *M. alfredi* and (B) *M. birostris* populations. Horizontal branch lengths reflect the amount of genetic drift that has occurred along each branch. Bootstrap support values for each node are indicated. Migration edges inferred using TreeMix are represented as arrows and coloured according to their migration weight. The scale bar reflects 10 times the average standard error of the entries in the sample covariance matrix. (C–E) Variation in individual multi-locus heterozygosity between (C) species and among populations of (D) *M. alfredi* and (E) *M. birostris*. Note that Y-axis scales differ for (D) and (E). Centre lines of boxplots reflect the median, bounds of the boxes extend from the first to the third quartiles, and upper and lower whiskers reflect the largest and smallest values but no further than 1.5 * the interquartile range from the hinge. Population abbreviations: AP = Australia Pacific, CHAG = Chagos, FIJI = Fiji, HAW = Hawaii, MAL = Maldives, SEY = Seychelles, MC = Mexico Caribbean, MP = Mexico Pacific, PERU = Peru, SA = South Africa, SL = Sri Lanka and PHI = the Philippines.

### Heterozygosity landscape across species and populations

To explore how patterns of population structure and colonisation are associated with genome-wide variation, we compared individual multi-locus heterozygosity between species and among populations. Strikingly, heterozygosity was on average over three times higher in *M. birostris* (mean = 0.10, min = 0.053, max = 0.12) than in *M. alfredi* (mean = 0.03, min = 0.01, max = 0.051), with every individual displaying a higher value than any *M. alfredi* (ß = 0.07, 95% CI = 0.06–0.07, *P* = <2.2 × 10^-16^, Figure 3C). This finding is in line with the patterns of population structure and historical splits we observed in each species. Variation in heterozygosity was also observed at a population level (Figure 3D–E). In *M. alfredi*, the Maldives and Australia Pacific had the highest levels of heterozygosity within each ocean basin, in line with these populations being first to split in the TreeMix analysis. Indian Ocean populations displayed higher overall heterozygosity than Pacific populations and had mean values that were overall similar. Interestingly, heterozygosity within Pacific populations declined steeply from west to east, with heterozygosity in Hawaiian individuals being around half that of the Australian animals (Figure 3D), in line with this population being last to split in the TreeMix analysis and displaying the highest amount of drift. In contrast, *M. birostris* populations displayed less extreme variation in heterozygosity, with mean values differing by less than 0.02 and having no clear geographical pattern (Figure 3E). Furthermore, variance within populations was an order of magnitude greater in *M. birostris* than in *M. alfredi*, and was particularly large in Sri Lanka, the Philippines, and Mexico Pacific populations.

## Discussion

Manta rays are iconic megafauna with cultural, socio-economic and ecological significance. Due to targeted and bycatch fisheries operating across their broad-ranging distributions, populations are declining worldwide. Elucidating levels of connectivity and genetic variation remains a crucial priority for conservation management. We use reduced-representation sequencing on a global set of samples and reveal striking differences in the population genetic landscape of two recently diverged manta ray species. By considering the influence of both contemporary and historical processes, our study provides a precautionary framework for assessing conservation units in widely distributed marine species.

We first demonstrate the presence of strong genetic differentiation in *M. alfredi* at a global and regional scale. From a total of six sampling locations, we found evidence for at least five genetically distinct, and by extension, demographically independent populations. Two of these were separated by a distance of ∼1,200 km, which is close to the maximum recorded movement in the species (Armstrong et al., 2019; Jaine et al., 2014), indicating that long distance migrations are likely rare. Indeed, contemporary gene flow was low – especially between geographically distant locations – with only a small proportion of individuals in any population being identified as first- or second-generation migrants. Furthermore, when gene flow was observed, it tended to be unidirectional. These results are in line with recent studies demonstrating population differentiation between Western Australia and Mozambique (Venables et al., 2021) and between Eastern Australia and New Caledonia (Lassauce et al., 2022), together highlighting how large ocean basins form significant barriers to dispersal in coastal elasmobranchs (Hirschfeld et al., 2021). High site-fidelity has been widely reported in *M. alfredi* based on tagging and mark-recapture studies (Braun et al., 2015; Deakos et al., 2011; Germanov et al., 2022; Jaine et al., 2014; Knochel, Hussey, et al., 2022; Peel et al., 2019; Setyawan et al., 2018). However, the degree of residency has been shown to vary, with movements rarely exceeding a few hundred kilometres in some locations (Braun et al., 2015; Deakos et al., 2011; Kessel et al., 2017; Knochel, Hussey, et al., 2022; Setyawan et al., 2018) yet reaching over 1,000 km in others (Armstrong et al., 2019; Germanov & Marshall, 2014). Our study presents a comparatively broad-scale analysis relevant for regional and global management planning. Further work on local patterns of population structure will shed light on the nuances and drivers of fine-scale movement patterns in this species (Whitney et al., 2023).

To explore the mechanism by which manta rays colonised their distribution, we reconstructed historical relationships and assessed levels of heterozygosity. In *M. alfredi*, we found strong evidence for an initial split between the Indian and Pacific Oceans followed by further separation within each locality. Furthermore, populations in the Pacific displayed a signal of declining heterozygosity from west to east, together suggesting that *M. alfredi* underwent a stepping-stone pattern of range expansion from an Indo-Pacific Ocean origin, involving opportunistic long-range movements and associated founder events. This is consistent with a recent observation of a pregnant *M. alfredi* individual at Cocos Island, Costa Rica (Arauz et al., 2019), almost 6,000 km east of the nearest confirmed sighting, and the first record of *M. alfredi* in the Eastern Pacific. Range expansion inherently impacts genetic variation, with a stepping-stone model of colonisation predicted to result in the strongest cumulative effect of founder events (Le Corre & Kremer, 1998). Among our sampled populations, Hawaii is the most geographically isolated, situated at the edge of the *M. alfredi* distribution. Interestingly, not only was Hawaii the most genetically differentiated from all populations in our study, but it displayed the longest external branch lengths in the TreeMix analysis and the lowest levels of heterozygosity. This could be suggestive of a single founder event by a small population. Genetic variation is fundamental for enabling populations to adapt in response to selection (Bonnet et al., 2022; Kardos et al., 2021; Lai et al., 2019). Our findings therefore expose how isolated *M. alfredi* populations at the periphery of their distribution may be intrinsically more vulnerable to changing environmental conditions and the genetic impacts of population decline.

In stark contrast to the patterns observed in *M. alfredi*, *M. birostris* displayed markedly higher levels of heterozygosity and only subtle genetic differentiation across ocean basins. Weak population structure is common in highly mobile marine species (Leslie & Morin, 2018; Nikolic et al., 2023; Vignaud et al., 2014), yet warrants careful interpretation, particularly considering management recommendations (Younger et al., 2017). On the one hand, these findings may be an indication of high contemporary gene flow and low natal philopatry, in line with the species’ occurrence at remote oceanic islands, tendency to range into sub-tropical habitats and lower overall re-sight rates than *M. alfredi* (Couturier et al., 2014; Harty et al., 2022; Rambahiniarison et al., 2023). To date, our understanding of the movement behaviour in *M. birostris* has largely been based on coastal aggregations of adult individuals over relatively short timeframes (Beale et al., 2019; Harty et al., 2022; Rohner et al., 2013; Stewart et al., 2016). Such studies have a tendency to capture seasonal migrations as opposed to dispersal events and may explain why only a handful of long-distance (∼1,000 km) movements have been recorded in the species (Andrzejaczek et al., 2021; Knochel, Cochran, et al., 2022). In an infinite island model, only a few migrants per generation are required to obscure strong population structure when *N_e_* is large (Wright, 1931) and therefore it is possible the patterns we observe translate to infrequent dispersal events. Furthermore, dispersal could be segregated by age and/or sex (McClain et al., 2022; Phillips et al., 2021), and may vary among individuals (Papastamatiou et al., 2013; Perryman et al., 2022; Thorburn et al., 2019). While challenging, there is benefit in extending future tagging efforts to transient individuals away from known aggregation sites (Garzon et al., 2023), as well as previously underrepresented age classes – such as juveniles – to capture what may be infrequent yet evolutionarily relevant movements.

An alternative explanation for the patterns we observe in *M. birostris* is that insufficient time has elapsed to reliably identify recent genetic divergence among localities. In contrast to *M. alfredi*, our TreeMix analysis indicated that *M. birostris* rapidly radiated from a large ancestral source, with only marginal genetic drift occurring between regions. This was further evidenced by substantially higher levels of genetic variation that differed little across sampling locations. In addition, little differentiation was observed between Mexico Pacific and Mexico Caribbean, two regions that have been geographically separated since the emergence of the Isthmus of Panama. These findings are consistent with a recent mark-resight analysis that estimated the population of *M. birostris* in coastal Ecuador to number at least 22,000 individuals (Harty et al., 2022). Large effective population sizes and high genetic variation increase the time taken for populations to diverge due to genetic drift (Bailleul et al., 2018; Taylor & Dizon, 1996; Wright, 1931). This is further compounded in species with long and overlapping generations (Hoffman et al., 2017) as is the case for manta rays (Dulvy et al., 2014). Taken together, genetic similarities among *M. birostris* localities may be partially confounded by recent shared ancestry and large effective population size.

On the basis of these considerations, we propose that a combination of large historical population size and contemporary gene flow have contributed to the comparatively high levels of heterozygosity and genetic homogeneity in *M. birostris*. The subtle population differentiation we observe between the Indian Ocean, South-East Asia and the Eastern Pacific is likely best explained by the geographic limits of dispersal as opposed to complete geographic isolation. Yet, unlike in *M. alfredi* where genetic clusters almost certainly reflect discrete demographic units relevant for conservation management, the extent to which genetic connectivity in *M. birostris* reflects demographic connectivity is less clear. For example, in extreme cases, the number of migrants required to eliminate signals of population structure will not be enough to demographically link populations, and more importantly, replenish those that have been depleted (Waples, 1998). Interestingly, while re-sight rates are typically lower in *M. birostris* than *M. alfredi*, demographic independence has been implicated in several mark-recapture studies where re-sightings follow predictable patterns (Beale et al., 2019; Cabral et al., 2023). Furthermore, a population genetic analysis based on *F_ST_* outliers uncovered allele frequency differences between two Mexican locations and Sri Lanka (Stewart et al., 2016), suggesting recent divergence against a background of ongoing gene flow. Taken together, we highlight the potential for further work investigating adaptive divergence between *M. birostris* populations and emphasise the need to combine molecular measures of connectivity with empirical demographic data in this species (Cayuela et al., 2018; Lowe & Allendorf, 2010; Younger et al., 2017).

### Conservation implications

The remarkable differences we observe in the population genetics of manta rays directly inform likely response to continued exploitation and respective conservation measures. At present, *M. alfredi* is among the most protected mobulid species worldwide, with some management frameworks in place at local, national, and international levels (Lawson et al., 2017; Stevens et al., 2018). Our findings of global population structure underline how local initiatives recognising populations as distinct management units will be most appropriate for this species. However, we also demonstrate the consequence of geographic isolation on genetic variation and reveal how *M. alfredi* likely faces a greater risk from local depletion. This is especially true for populations at the edge of the species range and in regions with high coastal fishing pressure. Prioritising these populations in conservation action plans and maintaining local connectivity will therefore be crucial for boosting resilience and preventing local extinction in this vulnerable species.

The implications of our findings for *M. birostris* are more nuanced. Despite detecting only subtle population genetic differentiation, we cannot rule out the possibility that historical processes and large effective population size are obscuring a higher degree of contemporary demographic separation. Together with studies reporting high site-fidelity and restricted movement patterns, our findings strongly suggests that local and national management action should be considered essential for protecting resident aggregations of *M. birostris*. Nevertheless, we expect that weak population structure and high genetic variation are simultaneously being driven by some degree of contemporary dispersal. Consequently, any fishing activity taking place along migratory corridors threatens to disrupt a mode of gene flow that may be fundamental for long-term resilience of the species. Similarly, although we have limited understanding of the number and distribution of breeding and nursery grounds (Knochel, Cochran, et al., 2022; Pate & Marshall, 2020; Stewart, Nuttall, et al., 2018), significant reduction of local stocks may impact long-term recruitment at oceanic and even global scales. We therefore emphasise the escalating need to improve the implementation of regional and international measures that seek to protect taxa in the high seas. Together with local scale management, appropriate evidence-based actions will contribute to maintaining large, connected and genetically diverse populations of manta rays into the future.

## Acknowledgements

We are grateful to the following people and organisations for their support sourcing and collecting tissue samples: Kerstin Forsberg, Jon Slayer, John Nevill, Dan Bowling, Heather Pacey, Karen Fuentes, Manta Caribbean Project, Annie Murray, Tam Sawers, Beth Taylor, Adam Thol’hath, Isabel Ender, Musa Mohamed, the Barefoot Collection, Four Seasons Resorts Maldives, Six Senses Laamu and all Manta Trust and LAMAVE staff, volunteers and field team. We are grateful to Helen Senn and Jennifer Kaden from the Royal Zoological Society of Scotland for support during library preparation. ddRAD sequencing was carried out by Edinburgh Genomics. Tissue sample collection in Seychelles was approved by the Seychelles Bureau of Standards, the Seychelles Ministry of Environment, Energy and Climate Change, and The University of Western Australia (RA/3/100/1480). Blue Resources Trust (BRT) would like to thank the Department of Wildlife Conservation and the Department of Fisheries and Aquatic Resources for enabling the fieldwork to be conducted in Sri Lanka. Sample collection in the Maldives was made possible thanks to the permission and support provided by the Ministry of Fisheries and Agriculture ([OTHR]30-D/INDIV/ 2016/220) and the Ministry of Environment and Energy (EPA/2016/PSR-M02). LAMAVE would like to thank the Bohol Environmental Management Office, the DA-Bureau of Fishery Region 7, the local communities and Mayors of Jagna and Balcayon (Bohol) and the LAMAVE field staff and volunteers including Joshua Rambahiniarison, Christina Pahang and Maita Verdote. Sampling in Mexico Caribbean was made possible thanks to CONAPESCA (PPF/DGOPA-126/16 and PPF/DGOPA/315717) and SEMARNAT (SGPA/DGVS/07618/15 and SGPA/DGVS/10067/17). We would like to acknowledge Richard Coleman for helpful comments on the manuscript. For the purpose of open access, the author has applied a Creative Commons Attribution (CC BY) licence to any Author Accepted Manuscript version arising from this submission.

## Author contributions

EH, JH, GC, MdB, SC, GMWS and RO conceived and designed the study. GMWS, AA, RB, MD, DF, NF, LRP, SP, AP, JDS and SW provided samples. JH and JK carried out laboratory work. EH analysed the data with input from JH. EH wrote the paper with input from all other co-authors.

## Data accessibility statement

Sequencing data have been deposited to the European Nucleotide Archive under study accession number <u>PRJEB66437</u>. Analysis code is available at https://github.com/elhumble/manta_pop_gen_2022.

## Benefit-sharing statement

A research collaboration was developed with scientists from regions providing genetic samples to advance the conservation of manta rays and their relatives through evidence based research, international collaboration and institutional capacity building. Data and analysis pipelines associated with this work have been shared on public databases for the benefit of this wider cause.

