## Supplementary Material for "Comparative population genomics of manta rays has global implications for management"

#### Table of contents

| | |
|---|---|
| Supplementary Methods | **Page 2** |
| Supplementary Figures | **Page 4** |
| Supplementary Tables | **Page 9** |
| References | **Page 11** |

### Supplementary Methods

##### DNA extraction and ddRAD library preparation

Whole genomic DNA was extracted using the Qiagen DNeasy Blood and Tissue Kit and quantified using a Qubit 3.0 Broad Range Assay. Double digest restriction-site associated DNA (ddRAD) libraries were prepared following (*1*) with modifications as described in (*2*). DNA was fragmented using restriction enzymes Sbf1 and Sph1 (NEB), and a unique pair of barcodes was ligated to the resulting fragments. Samples were then pooled into three libraries and size-selected between 400 and 700 bp using gel electrophoresis. Samples from different species and populations were multiplexed together within the libraries to minimise the risk of batch effects. Each library was then PCR amplified to incorporate adapter sequences. The resulting libraries were 125 bp paired-end sequenced on an Illumina HiSeq.

##### Data processing and quality control

Sequencing reads were demultiplexed using process_radtags in Stacks v2.54 (*3*) and assessed for quality using FastQC. A total of 12 individuals were excluded from subsequent analysis due to low read numbers (Figure S6). We then followed the procedure outlined in (*4*) for *de novo* assembly and genotyping using Stacks. First, we optimised the *de novo* assembly parameters by determining the combination that yielded the highest number of polymorphic loci in at least 80% of individuals. For this, -*m* was set to 3, -*M* was assigned values ranging from 1 to 9 and -*n* was set as equal to -*M* or one unit greater to account for fixed polymorphisms in either *Mobula alfredi* or *Mobula birostris*. This procedure was run on a subset of two individuals with the highest number of reads from each location across both species. The optimal combination of parameters ($m$ = 3, $M$ = 3, $n$ = 4, Figure S7) was then used to analyse the complete dataset. When running the cstacks module, only a subset of samples were used to generate the catalog to reduce both noise and computation time. For this, up to 10 individuals with the highest depth of coverage were selected from each location across both species, discounting those with a depth of coverage less than 25.

##### SNP filtering

We applied a set of initial filters to the raw genotypes using VCFtools v1.13 in order to remove low quality sites and individuals. For this, genotypes with a depth of coverage less than 6, SNPs called in less than 40% individuals and individuals with more than 45% missing data were removed. We then filtered SNPs more stringently by removing those called in less than 60% of individuals and with a depth of coverage less than 25 or greater than 376 (equivalent to 95% quantiles for mean depth of coverage). This resulted in a set of 81,166 sites called in 181 individuals with an average coverage of 137.4, that was used to generate three final datasets for subsequent analyses: two high-quality species-specific datasets; and one high-quality dataset comprising both species. For the species-specific datasets, we first extracted either *M. birostris* or *M. alfredi* individuals. Second, we identified and removed related individuals using the procedure described below. Third, SNPs were pruned for linkage disequilibrium using the -indep function in PLINK v1.9 with a sliding window of 50 SNPs, a step size of 5 SNPs and variance inflation threshold of 2. Finally, SNPs with a minor allele count of less than 3 and a genotyping rate less than 90% were removed using PLINK. This left a total of 1,553 SNPs in 91 *M. alfredi* individuals, and 6,278 SNPs in 82 *M. birostris* individuals. These datasets were used for analysis of population structure and contemporary migration. For the dataset comprising both species, we first removed individuals identified as being related, and then removed SNPs with a minor allele count of less than 3 and a genotyping rate less than 90%. This left a total of 15,312 SNPs called in 91 *M. alfredi* and 82 *M. birostris*. This dataset was used for the TreeMix and heterozygosity analysis.

**Related individuals**

To estimate relatedness among individuals, we first pruned species-specific SNP datasets for linkage disequilibrium using the --indep function in PLINK, a sliding window of 50 SNPs, a step size of 5 and a variance inflation factor threshold of 2. We also removed SNPs that deviated significantly from HWE with a p-value threshold of 0.001 and with a minor allele frequency < 0.3. We then estimated KING, R0 and R1 coefficients (*5*) using NgsRelate v2 (*6*). These statistics are based on genome-wide patterns of identity by state sharing between two individuals. Four pairs of individuals in *M. alfredi* (1352 / 1355, 1352 / 1356, 1355 / 1356 and 1293 / 1291) and five in *M. birostris* (1061 / 1060, 0728 / 0729, 1121 / 1122, 1077 / 1078 and 1139 / 1137) fell above the KING-robust kinship threshold for first-degree relatives (Figure S8). Each pairing comprised individuals from the same geographic location. The following individuals were removed from subsequent analysis: 1355, 1356, 1293, 0728, 1077, 1061, 1121 and 1137. The remaining individuals displayed no sign of close relatedness (*M. alfredi* mean pairwise relatedness = 0.06, *M. birostris* mean pairwise relatedness = 0.009).

#### Supplementary Figures

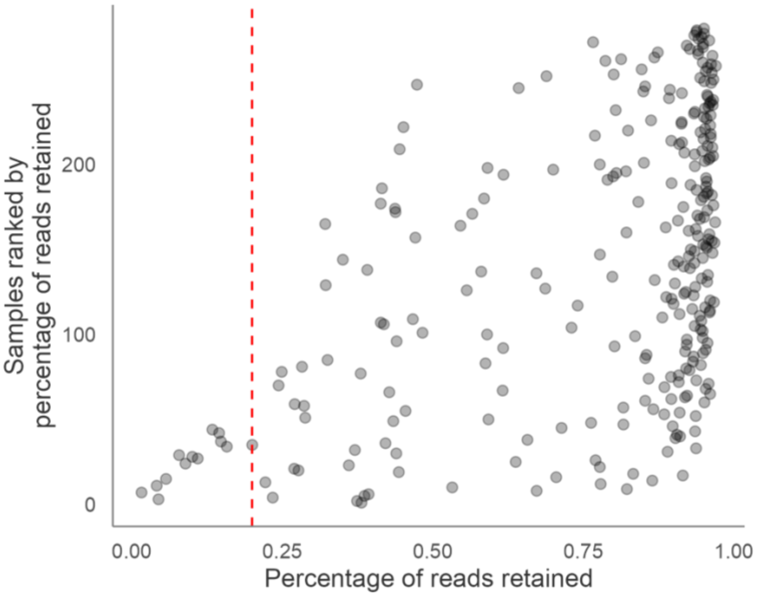


**Figure S1:** Distribution of the percentage of reads retained for each sample, ranked in ascending order. The dashed red line shows the percentage of reads retained below which samples were discarded (20%).

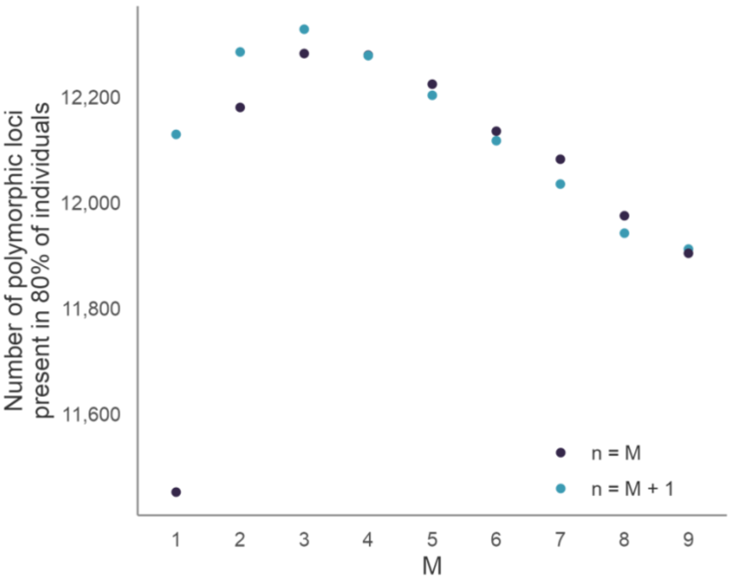


**Figure S2:** Change in the number of polymorphic loci present in at least 80% of the samples for increasing values of the *M* and *n* assembly parameter in STACKS. The light and dark blue points correspond to assemblies performed by setting *n* = *M* and n = M + 1 respectively. The number of assembled loci reached its peak at *M* = 3 and *n* = 4.

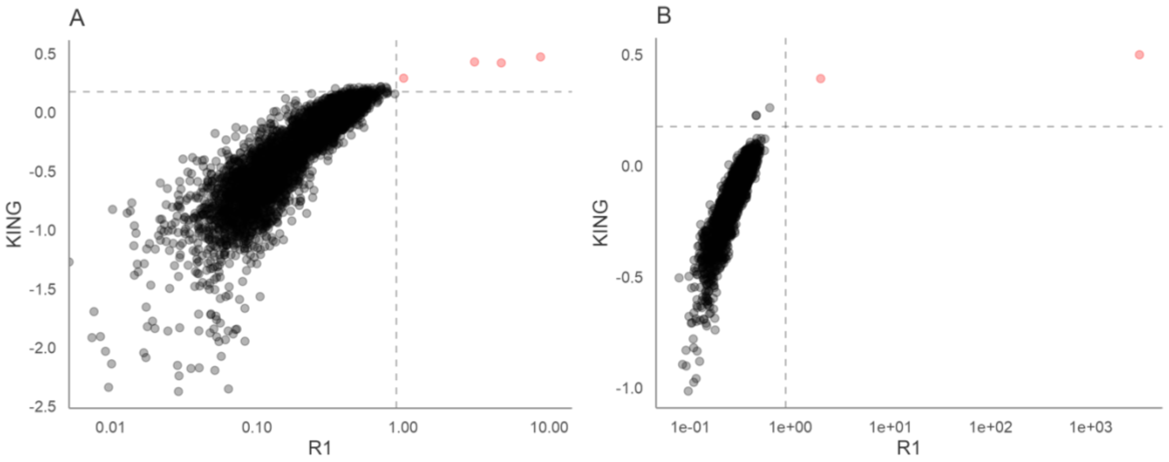


**Figure S3:** Empirical relatedness for all individual pairwise comparisons for (A) *M. alfredi* and (B) *M. birostris*. R1 coefficients are plotted against KING-robust kinship coefficients. The dashed line represents the KING-robust kinship and R1 threshold for first degree relatives. Four and two pairs of individuals fell above this threshold for (A) *M. alfredi* and (B) *M. birostris* respectively, and are colour coded in red.

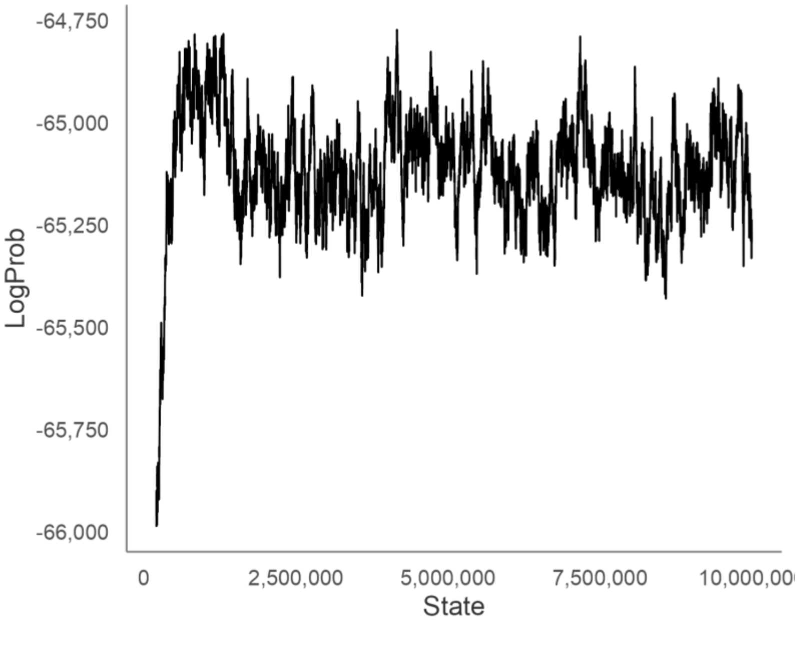


**Figure S4:** Trace profile of log probabilities for each iteration of the BayesAss MCMC analysis implemented in BA3-SNPs (-seed 10). BayesAss was run with 10,000,000 iterations, a burn-in of 1,000,000 and a sampling interval of 1000. The log probability increases steeply during the burn-in phase, after which it oscillate around a plateau indicating chain convergence.

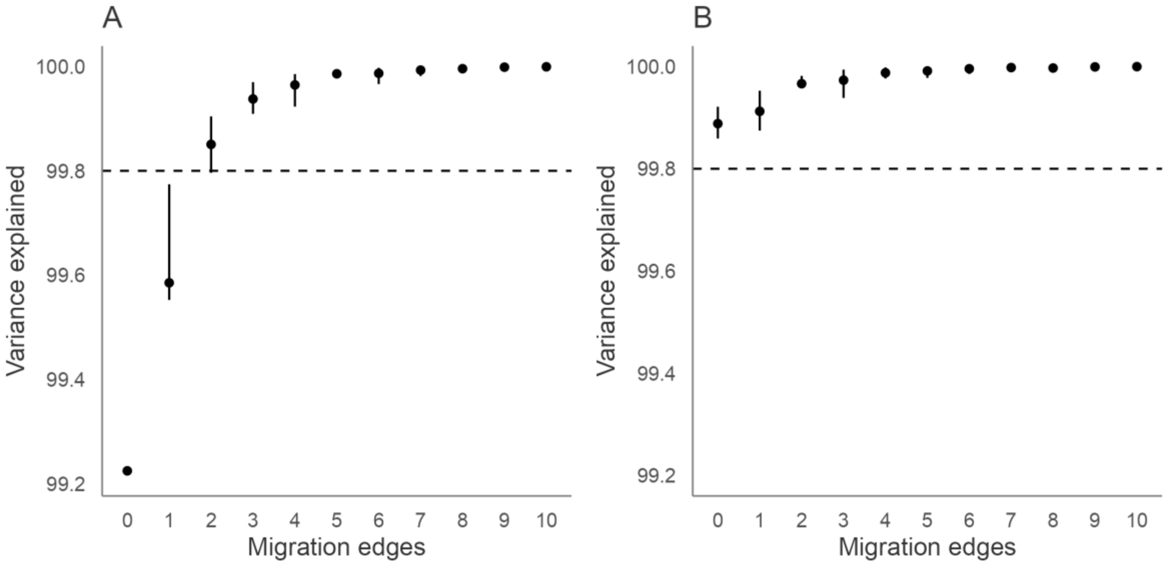


**Figure S5:** Variance explained in the TreeMix maximum likelihood models with increasing number of migration edges for (A) *Mobula alfredi* and (B) *Mobula birostris*. Points represent mean values and error bars represent minimum and maximum values for 10 initial runs of TreeMix per migration edge value.

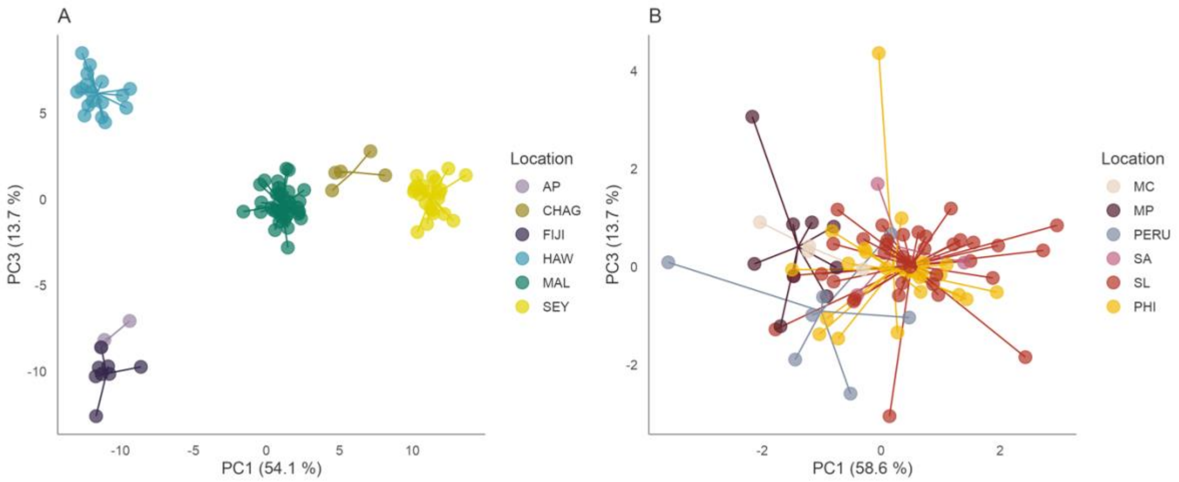


**Figure S6:** Scatterplots showing individual variation in principal components (PC) one and three derived from discriminant analysis of principal components analysis for (A) *M. alfredi* and (B) *M. birostris* individuals. The amount of variance explained by each PC is shown in parentheses. Population abbreviations: AP = Australia Pacific, CHAG = Chagos, FIJI = Fiji, HAW = Hawaii, MAL = Maldives, SEY = Seychelles, MC = Mexico Caribbean, MP = Mexico Pacific, PERU = Peru, SA = South Africa, SL = Sri Lanka and PHI = the Philippines.

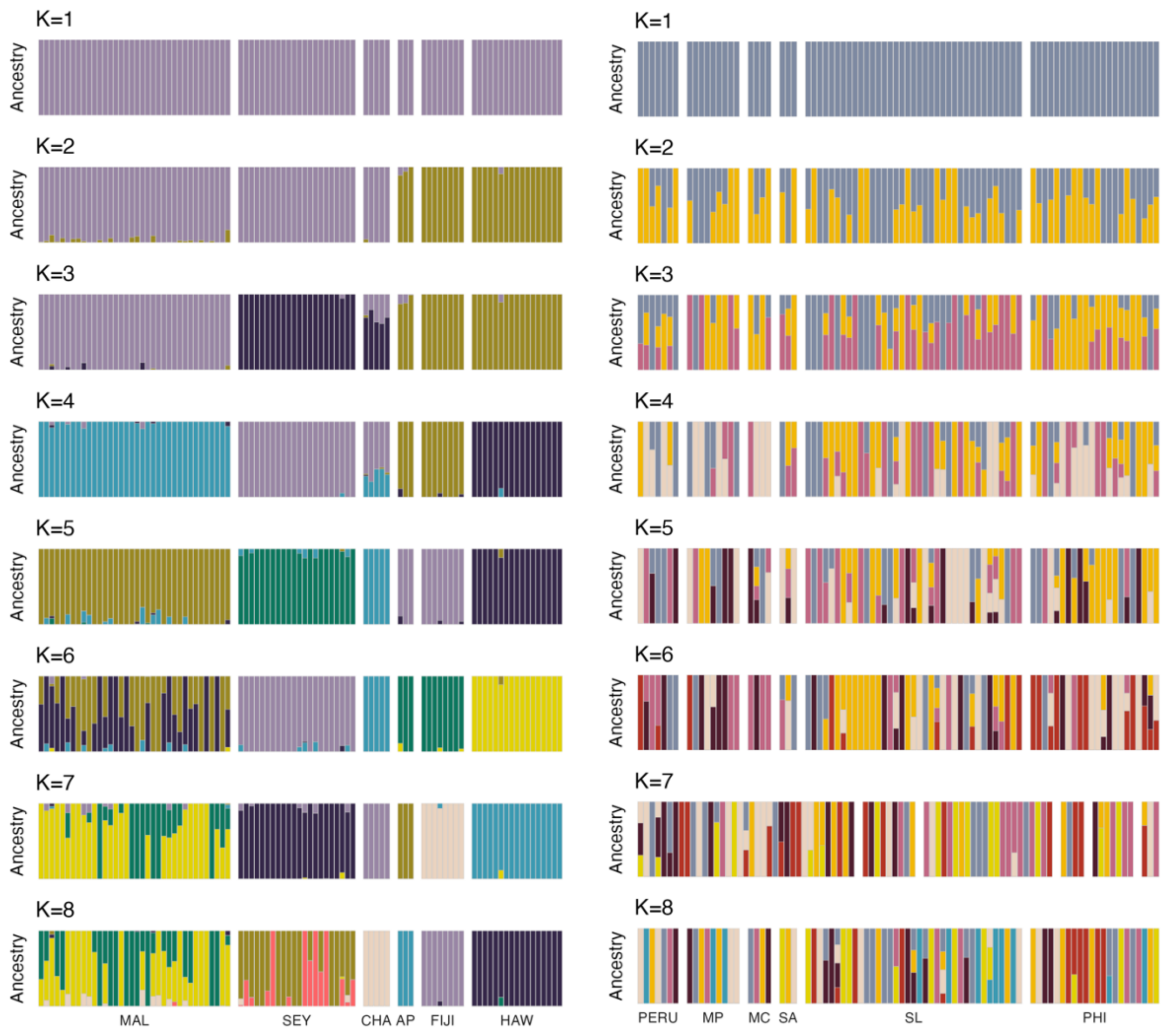


**Figure S7:** Ancestry proportions for each of the (left) 91 *M. alfredi* and (right) 82 *M. birostris* individuals inferred using ADMIXTURE for $K$ = 1 to $K$ = 8. Population abbreviations: AP = Australia Pacific, CHAG = Chagos, FIJI = Fiji, HAW = Hawaii, MAL = Maldives, SEY = Seychelles, MC = Mexico Caribbean, MP = Mexico Pacific, PERU = Peru, SA = South Africa, SL = Sri Lanka and PHI = the Philippines.

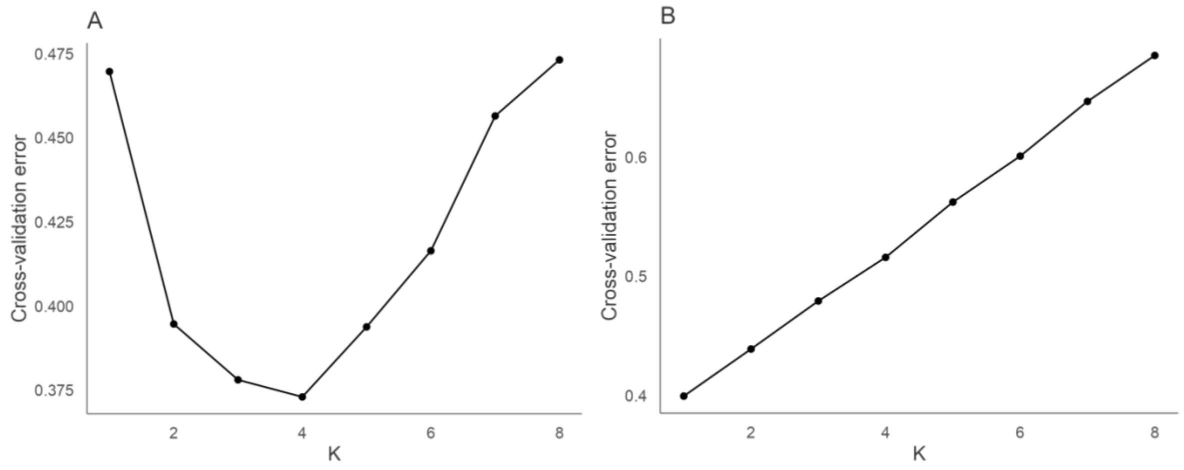


**Figure S8:** ADMIXTURE cross-validation error for $K = 1$ through to $K = 8$ for both (A) *M. alfredi* and (B) M*. birostris*.

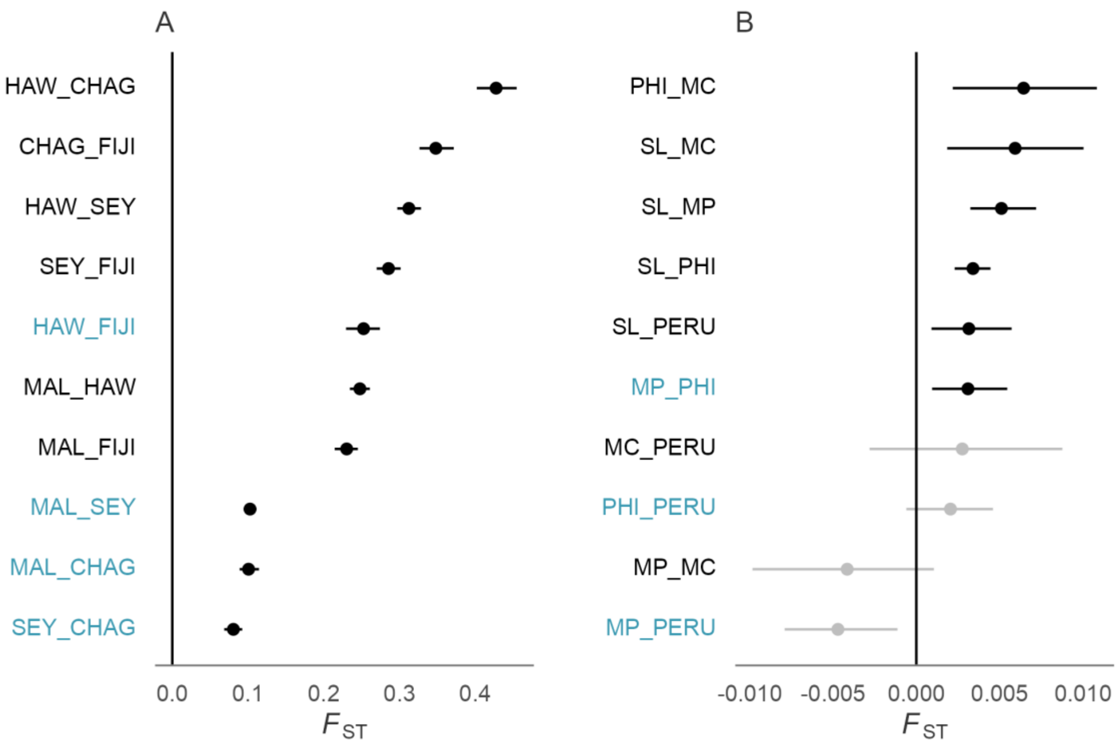


**Figure S9:** $F_{ST}$ estimates and confidence intervals for all pairwise population comparisons in (A) *M. alfredi* and (B) *M. birostris*. Significant comparisons are indicated in black and non-significant comparisons are indicated in grey. Population abbreviations: CHAG = Chagos, FIJI = Fiji, HAW = Hawaii, MAL = Maldives, SEY = Seychelles, MC = Mexico Caribbean, MP = Mexico Pacific, PERU = Peru, SL = Sri Lanka and PHI = the Philippines. Within ocean comparisons are noted blue, between ocean comparisons are in black.

### Supplementary Tables

**Table S1.** Genetic diversity summary statistics calculated from species and population datasets: Sample size (*N*), allelic richness (*Ar*), observed heterozygosity ($H_O$), expected heterozygosity ($H_E$), unbiased expected heterozygosity ($H_E$ unbiased), inbreeding coefficient ($F_{IS}$), *p*-values from chi-square test for goodness-of-fit to Hardy–Weinberg equilibrium (HWE), significance tests of homozygote and heterozygote deficiency. 95% confidence limits are shown in brackets.

| | *N* | *Ar* | $H_O$ | $H_E$ | $H_E$ unbiased | $F_{IS}$ | Test global HWE | Test HWE (homozygote deficiency | Test HWE (heterozygote deficiency) |
|---|---|---|---|---|---|---|---|---|---|
| **Species** | | | | | | | | | |
| *Mobula birostris* | 82 | 1.42 [1.39–1.45] | 0.101 | 0.115 | 0.115 | 0.112 [0.08–0.118] | 1.000 | 1.000 | 0.075 |
| *Mobula alfredi* | 91 | 1.15 [1.13–1.17] | 0.033 | 0.05 | 0.05 | 0.221 [0.204–0.255] | 0.000 | 1.000 | 0.000 |
| ***Mobula alfredi*** | | | | | | | | | |
| Maldives | 36 | 1.42 [1.33–1.46] | 0.198 | 0.199 | 0.202 | 0.012 [-0.027–0.023] | 1.000 | 1.000 | 1.000 |
| Seychelles | 22 | 1.42 [1.34–1.46] | 0.196 | 0.197 | 0.202 | 0.011 [-0.052–0.028] | 1.000 | 1.000 | 1.000 |
| Chagos | 5 | 1.33 [01.17–1.41] | 0.169 | 0.165 | 0.186 | -0.045 [-0.395–-0.033] | 1.000 | 1.000 | 1.000 |
| Fiji | 8 | 1.25 [1.14–1.29] | 0.13 | 0.121 | 0.13 | -0.066 [-0.225–-0.075] | 1.000 | 1.000 | 1.000 |
| Australia Pacific | 3 | 1.3 [1.13–1.37] | 0.169 | 0.137 | 0.165 | -0.231 [-1–-0.231] | 1.000 | 1.000 | 1.000 |
| Hawaii | 17 | 1.16 [1.11–1.19] | 0.074 | 0.078 | 0.08 | 0.048 [-0.046–0.067] | 1.000 | 1.000 | 1.000 |
| ***Mobula birostris*** | | | | | | | | | |
| Sri Lanka | 37 | 1.36 [1.26–1.40] | 0.149 | 0.166 | 0.168 | 0.081 [0.034–0.093] | 1.000 | 1.000 | 1.000 |
| The Philippines | 22 | 1.36 [1.26–1.40] | 0.151 | 0.165 | 0.169 | 0.065 [0.01–0.071] | 1.000 | 1.000 | 1.000 |
| South Africa | 3 | 1.30 [1.09–1.39] | 0.165 | 0.141 | 0.172 | -0.179 [-1–0.179] | 1.000 | 1.000 | 1.000 |
| Mexico Caribbean | 4 | 1.31 [1.10–1.39] | 0.157 | 0.146 | 0.168 | -0.095 -0.373–-0.095] | 1.000 | 1.000 | 1.000 |
| Peru | 7 | 1.35 [1.28–1.39] | 0.152 | 0.155 | 0.167 | -0.008 [-0.166–-0.014] | 1.000 | 1.000 | 1.000 |
| Mexico Pacific | 9 | 1.33 [1.12–1.40] | 0.137 | 0.159 | 0.169 | 0.09 [-0.044–0.107] | 1.000 | 1.000 | 1.000 |

**Table S2.** Contemporary migrates rates inferred using BA3-SNPs and BayesAss between locations of *M. alfredi*. Values in bold indicate significant migration rates where 95% credible sets (mean migration rate ± 1.96 x mean standard deviation) did not overlap zero. Values along the diagonal indicate the mean proportion of non-migrants within a location.

| | Sink | | | | | |
|---|---|---|---|---|---|---|
| Source | Australia Pacific | Chagos | Fiji | Hawaii | Maldives | Seychelles |
| Australia Pacific | 0.7039 | 0.0302 | 0.0238 | 0.0145 | 0.0079 | 0.0118 |
| Chagos | 0.0368 | 0.7276 | 0.0239 | 0.0145 | 0.0112 | 0.0120 |
| Fiji | **0.1478** | 0.0301 | 0.8808 | 0.0146 | 0.0080 | 0.0120 |
| Hawaii | 0.0370 | 0.0302 | 0.0239 | 0.9275 | 0.0080 | 0.0119 |
| Maldives | 0.0373 | 0.0304 | 0.0238 | 0.0145 | 0.9571 | 0.0119 |
| Seychelles | 0.0373 | **0.1515** | 0.0239 | 0.0144 | 0.0079 | 0.9403 |
